# Potential of fungicides, botanicals and biocontrol agents to induce physio-biochemical tolerance on *Curcuma longa* impaired by *Colletotrichum gloeosporioides*

**DOI:** 10.1101/2021.02.11.430813

**Authors:** Nasreen Musheer, Shabbir Ashraf, Arshi Jamil

**Author notes:** Corresponding Author: Nasreen Musheer.

## Abstract

Necrotic leaf spot of *Curcuma longa* (turmeric) limits the chief physio-biochemical activity for maintaining the plant health and productivity. In the present study, polyhouse and open field trials were conducted to estimate the pathogenicity of *C. gloeosporioides* on turmeric and to evaluate the foliar efficiency of propiconazole @ RD and copper oxychloride, extracts of *A. indica, A. sativum* and *O. sanctum* @ 40%, and culture filtrates of *T. viride, T. harzianum* and *T. virens @* 4×10^6^ cfu/ml in inducing physio-biochemical tolerance of pathogen inoculated and non-inoculated plants. In both the trials, these three agents yielded the highest efficiency to enhance the physio-biochemical traits. The induced physio-biochemical tolerance in treated turmeric plants showed variation in the elevation of plant health and immunity in response to pathogen aggressiveness or disease severity. However, phytophenol content was quite higher in infected plants than non-infected plants due to initiation of defense reaction in response of pathogenic elicitors. Thus, the present study demonstrated the novelty of physio-biochemical tolerance induction on turmeric plants by using fungicides, biocontrol agents and phytoextracts.

**Highlights:**

- Foliar treatments improve desirable plant physio-biochemical traits against pathogen.
- Physio-biochemical variation induces the innate plant defense system.
- High phytophenol accumulation counteracts the pathogenic stress.
- Turmeric plant’s health and yield enhance by the reduction of disease intensity.

## 1. Introduction

Turmeric (*Curcuma longa* L.) is one of the most important annual, monocotyledonous rhizomatous crop (Vasala et al. 2012). The demand of turmeric cultivation at national and international level is increasing profusely day by day in order to support the green health remedies and culinary purposes. Rhizome is the chief source of reserving essential bio-secondary metabolites including alkaloids, glycosides, coumarins, flavonoids, steroids, corticosteroids, essential oils, etc. (Amalraj et al. 2016). India is one of the known leading countries in production, consumption and export services of turmeric (Anuradha et al. 2018). Successful cultivation of turmeric in different states of India was found to be suppressed by abiotic and biotic stress conditions (Singh et al. 2013; Anandaraj et al. 2014). Amongst the diseases, leaf spot is a severe fungal disease caused by hemibiotrophic pathogen, *Colletotrichum gloeosporioides* (Panz. and Sacc.); this decease leads to significant yield loss (>50%) (Hudge and Ghugul 2010). The management of turmeric leaf spot disease caused by pathogen *Collettorichum* spp. by using fungicides, biocontrol agents and botanicals extracts, and their influence on the physio-biochemical activity and growth of host plants have been focus of intensive research. The disease symptoms of turmeric are characterized by producing brown, necrotic and sunken lesions of ashy center and surrounded by yellow halo or sometime numerous black dots like a structure called ‘acervuli’ formed in a concentric manner on leaves during September and October months (Ramakrishnan, 1954; Adhipath et al. 2013). The prevailing atmospheric conditions are favored by continuous rain, high humidity 80-90% and optimum temperature, which cause great loss in yield of rhizome up to 62.5% (Mishra and Pandey 2015).

Plants’ interaction with pathogen is recognized by microbial molecules called elicitors, which cause modification in host-physio-biochemical functions. Elicitors are classified as pathogen-associated molecular patterns that induce pattern-triggered immunity (PTI) in host plant by pattern recognition receptors and effector-associated virulent pathogens that contribute to effector-triggered immunity (ETI) by encoding R genes (Jones and Dangl, 2006; Smakowska-Luzan et al. 2018). Fungal-effectors of susceptible host plant promotes pathogenesis by interfering in both PTI and ETI (Tan et al. 2010; He et al. 2020). Infection caused by pathogen *Colletotrichum* sp. limits the photosynthesis or other physiological process of host plants (Berger et al. 2007 a, b; Guerra et al. 2014; Dallagnol et al. 2015). The deterioration of leaf cuticle and chloroplast cells is associated with the decline of photo-pigments concentration, viability as well as numbers of stomata with progress of plant-fungus parasitic interaction (Resende et al. 2012). Therefore, stomatal closure limited the absorption of light and disturbed the water equilibrium in plant, thus suppressing both transpiration rate and stomatal conductance during infection (Lobato et al. 2009). The diffusion of CO_2_ from surrounding atmosphere into plants increases the synthesis of new carbohydrate molecules serves as source for biomass enhancement (Van der Kooi et al. 2016; Thompson et al. 2017). *Colletotrichum truncatum* anthracnose disease of soybean affects the physiological performance and causes great loss in the yield (Dias et al. 2018).

Previous studies have suggested an integrated approach for disease management by using cultural, mechanical, biological and chemical controls (Wharton and Dieguez-Uribeondo, 2004; Agrios 2005). Several parameters were studied under phyto-physio-biochemical tolerance, such as assimilation rate, transpiration rate, photosynthesis, stomatal conductance and bioaccumulation of chlorophyll, carotenoid, phenol of leaf tissues and curcumin or oleoresin of rhizome tissues. Improved physio-biochemical functioning of plants is proportionally related to treatments. A major threat of physio-biochemical process is *C. gloeosporioides* that reduces the tolerance level in plants (Castro et al. 2016). The localized infection in plant tissue upregulates defense genes to PR-proteins (salicyclic acid), enzymes (Chitinases and β1,3-glucannases) and transcription factors (expression of genes by interacting with cis-elements present in the promoter region), which are known to activate physio-biochemical tolerance in plants against pathogen (Koch et al. 2016; Israel Pagan and Fernando 2018; Ethan et al. 2018). Accordingly, genes expression of proteins and enzymes (particularly involved in glycolysis and Krebs cycle) provides the adequate energy to enhance various metabolic processes of the plants to increase tolerance level against stress (Kosova et al. 2018; Xing et al. 2019).

Mishra and Pandey (2015) reported that foliar application of propiconazole (0.1%) was significantly superior in reducing the disease intensity of leaf spot (27.61 PDI) and increasing fresh rhizome yield (ranged from 33.96 - 34.33 t ha^-1^ over the control (28.17t ha^-1^). Biocontrol agents enhance the desirable growth promoting characteristics and physio-biochemical changes in plants, which increases plant tolerance to pathogen by triggering plant defense system of induced systemic resistance (ISR) or systemic acquired resistance (SAR). *Trichoderma* species offer various mechanisms such as mycoparasitism and secretes (diffusible, volatile and non-volatile antifungal secondary metabolites) against both soil-borne (e.g. *R. solani* and *Sclerotium rolfsii)* and foliar pathogens like *Botrytis cinerea* (Lopez-Mondejar et al. 2011; Li et al. 2016). Jagtap et al. (2013) reported that high efficacy of *Trichoderma harzianum, Trichoderma viride, Gliocladium* spp., *Trichoderma koningii* and *Pseudomonas fluorescens* in the reduction of mycelial growth of *C. capsici* caused leaf spot disease of turmeric *in-vitro* conditions. Hung et al. (2013) also proved that the use of *Trichoderma viride* volatile organic compounds (VOCs) on *Arabidopsis thaliana* enhanced the biomass of plants and chlorophyll concentration. Many Phytoextracts possess antifungal properties against the phytopathogens, which could be used commercially without causing any toxic residual effect on ecosystem (Kumar et al. 2007). The extract of azadirachtin was suppressed 89.25% mycelial growth of *C. gloeosporioides* (Hegde et al. 2014). Similarly, Jagtap et al. (2013) and Musheer et al. (2019) studied the foliar efficacy of fungicides, biocontrol agents and botanical extracts to enhance the growth and yield traits against the *C. gloeosporioides*.

While the use of fungicides, biocontrol agents and phytoextracts to enhance the growth yield of turmeric plant has been widely studied, there is lack of attention on their application to induce physio-biochemical tolerance on plants. Producing stress-tolerant variants of plants against biotic constrains by modifying the physio-biochemical traits has emerged as an efficient, viable and cost-effective approach. Accordingly, the present work has focused on the foliar application of fungicides, biocontrol agents and phytoextracts in the improvement of physio-biochemical activity of turmeric under pathogenic stress. Ployhouse and open-field experiments were conducted on turmeric plants for a period of four years (2015-19), and the data were analyzed by using statistical tools.

## 2 Materials and Methods

### 2.1. Isolation and identification of pathogen and biocontrol agents

The isolation of *Colletotrichum* sp. was done on Colletotrichum specific Mathur’s medium modified (peptone, magnesium sulphate, heptahydrate, potassium dihydrogen phosphate, sucrose and agar) by placing the infected pieces of turmeric leaves that showed the typical symptoms of necrotic or brown lesion in different shapes and sizes. Inoculated plates were allowed to incubate at 27±1° C till the appearance of mycelial growth and then purified by using hyphal tip culture method. *Trichoderma* was isolated from rhizosphere soil around the healthy turmeric plants by using *Trichoderma* selective medium (TSM). The Potato dextrose agar (PDA) medium was used to maintain the pure culture of pathogen and biocontrol agents.

#### 2.1.1. Morphological characterization

Identification was done on the basis of morphological characteristics like growth rate, pattern and colours of cultural growth on plate, whereas microscopic structures such as conidiophores, conidia, phialides or mycelium were visualized under high resolution (10X objective×10X ocular) compound binocular microscope.

#### 2.1.2. Molecular characterization

Morphologically identified isolates of *Colletotrichum* sp. and *Trichoderma* species were assessed for PCR amplification of 18s rRNA-internal transcribed spacer (ITS) of ribosomal RNA region by using universal primers ITS1 5’ (TCCGTAGGTGAACCTGCGG) 3’ and ITS4 5’ (TCC TCCGCTTATTGATATGC) 3’. The PCR amplified products were of approximately 590bp size, which were obtained from Macrogen® Incorporation, South Korea. The purified nucleotide sequence was run in the nucleotide Basic Local Alignment Search Tool (BLAST) of the National Centre for Biotechnology Information (NCBI), in order to match with the available standard nucleotide database for exact species confirmation. The analyzed nucleotide sequence of pathogen and biocontrol agents had to be submitted to the GenBank of NCBI to acquire the specific accession number.

### 2.2. Preparation of culture filtrates

The culture filtrates of *Trichoderma* spp. and *Colletotrichum* sp. were prepared in nutrients broths (beef extract, yeast extract, peptone and sodium chloride) contained by 250ml Erlenmeyer flasks. Excised 5mm segment from the periphery of seven days old pure culture plates has transferred aseptically in to the broth and incubated at room temperature (27±1^0^C) till the appearance of mycelial disc on the surface. Subsequently, the culture broth was filtered through double-layer cheese cloth and the spore suspension was then centrifuged at 5000rpm at 28°Cfor 5 minutes. The conidial mass was collected in sediments and was re-suspended in sterile distilled water to maintain homogeneity of the suspension. The spore density per ml was measured by using haemocytometer.

### 2.3. Preparation of phytoextracts

Crude extraction of *Azadirecta indica* (neem leaf), *Ocimum sanctum* (tulsi leaf) and *Allium sativum* (garlic bulb) in distilled water (1:1w/v) were accomplished through a double-layered muslin cloth and Whatman No. l filter paper.

### 2.4. Pot and field experiments

The experiments were conducted for four years: pathogenicity test was conducted during the first year (2015-16) followed by polyhouse (2016-17) and field (2017-19) experiments in the department of Plant Protection, Aligarh Muslim University Aligarh (India) at 27°52.887′N latitude and 78°4.4784′E longitude with elevation of 189 m above sea level.

The turmeric var. sogandham was grown in earthen pots of 20cm×30cm size filled with homogenous mixture of sterilized soil, decomposed farmyard manure and vermicompost in 2:1:1 ratio. Microplots of size 1.5×2m^2^ were prepared in a field area of 180m^2^. The healthy rhizome @ one rhizome/pot or five holes per ridges was sown in the third week of May (in 2015, 2016 2017, 2018 and 2019) when the pre-monsoon shower is available to promote the budding/germination. Each microplot was maintained at interplant spacing of 40cm and inter-ridge spacing of 30cm with total number of five ridges and five holes per ridge, to avoid the moisture accumulation that is conducive for disease development. The planted pots were maintained under controlled environment conditions of polyhouse where they received favorable ambient conditions (temperature: 28-30ºC, and relative humidity (80-90%)) for the efficient growth of turmeric. However, field soil of sandy loamy texture with pH 6.5 was fertilized at recommended dose (60 kg N: 60 kg P2O5: 120 kg K2O), FYM @ 4t/ha and neem cake @ 2t/ha applied as basal dressing and ploughed 2-3 times based on soil physiochemical results through Agriculture farmer welfare Ministry of Government of India, Aligarh. Chemical fertilizers half dose of N, full dose of P and half dose of K were applied before planting thereafter remaining half doses of both N and *K* were used at 45 and 90 days after planting (DAP) per hectare. (Roy and Hore 2012; Shamrao et al. 2013; Modupeola 2015). The pots and microplots were irrigated regularly with tap water till the harvest.

### 2.4. Pathogenicity test (2015-2016)

The virulence effect of *C. gloeosporioides* on turmeric var. sogandham was measured at four different doses of pathogen inoculum under pot condition, during 2015-2016. Fifty ml spore suspension at conidial loads of 1×10^6^, 2×10^6^, 3×10^6^ and 4×10^6^ cfu/ml were adjusted via haemocytometer. These spore suspensions were sprayed on pricked leaves of turmeric at 3-4 leaf stage after three months of planting under the polyhouse conditions. The inoculated pots were incubated for 15 days or till the appearance of symptoms inside the poly house where they received the favorable ambient conditions (28±2ºC temperature and 90% relative humidity) essential for the disease development. Three replicates of each dose were maintained in completely randomized block design.

### 2.5. Foliar treatment: Polyhouse and field trials (2016-2019)

The subsequent active season (2016-2017) of turmeric cultivation was selected for the screening of fungicides, botanicals and biocontrol agents against the pre inoculated pathogen @ 3×10^6^ cfu/ml and non-inoculated stage of the plants in pots. The pathogen @ 3×10^6^ cfu/ml was found to cause reasonable damage to the plants. Therefore, this dose of pathogen inoculation was used for the evaluation of three foliar sprays of fungicides, botanical extracts and biocontrol agents at 45 days interval after planting for the enhancement of both physiological and biochemical characteristics of the plant, which cause higher biomass under control environment conditions of polyhouse. Subsequently, field trial was done during 2017-2019 to screen each treatment efficiency against the natural occurrence of leaf spot disease under varied environmental conditions. Two fungicides *viz*. propiconazole and carbendazim12%+mancozeb63% were used @ lower than recommended dose (LRD), recommended dose (RD) and higher than recommended dose (HRD); botanical extracts of *A, indica*; *A. sativum* and *Sanctum* @40%; and culture filtrate of *T. viride, T. harzianum* and *T. virens* @ 4×10^6^ cfu/ml. Each treatment was replicated in six pots and three microplots in a randomized block design (RBD) manner.

The brown spots severity on leaves was recorded at 0-9 disease grade scale (0= no symptom; 1= less than1% leaf area covered with brown spot; 3= 1-10% leaf area covered with lesions; 5= 11-20% leaf covered with brown lesions; 7= 21-50% brown lesions; 9= ≥51% leaf area infected). The percent severity index (PSI) was calculated by Eq. (1):

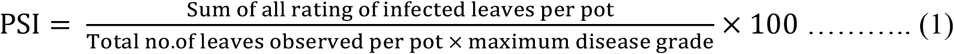

### 2.6 Assessment of leaf area

Leaf area (LA) was calculated at 180 DAP by selecting five leafs per plant randomly. The length and breadth were measured by using leaf area constant K = 0.6454 (Rao et al., 1994).

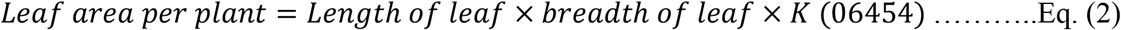

### 2.7. Assessment of physiological parameters

Transpiration rate (T_N_) net photosynthetic rate (P_N_), and stomatal conductance (gs) were estimated at 150 and 180 days after planting (DAP). The tip of the fresh, fully expanded leaves was placed in a portable Infra-Red Gas Analyzer of photosynthetic system (IRGA) (LICOR 6400, Lincoln, Nebraska, USA). The gas analyzer was calibrated at zero for every half an hour during the measurement period and the data of each treatment were measured thrice.

### 2.8. Assessment of phytochemical contents

The leaf samples of 150 and 180 DAP were collected for the quantitative assessment of essential leaf constituents, and the rhizome constituents were measured after completion of the harvesting period by using UV/VIS-spectrophotometer (UV-Pharma Spec 1600, Shimadzu, Japan).

#### 2.8.1. Extraction of chlorophyll and carotenoids contents

The quantitative analysis of specific photosynthetic components such as total chlorophyll and carotenoid in milligram per gram of fresh leaves tissues were analyzed by following the technique of Musheer et al. (2019). One gram of fresh leaf was crushed in 5 ml of 99.9% (v/v) acetone using mortar and pestle and the suspension was filtered through Whatman filter paper number 1. The final filtrate volume was made up to 10 ml by adding acetone, followed by centrifugation at 15,000 rpm for 10 min at 10 °C. Before recording the new absorbance reading at a particular wavelength, the absorbance reading must be calibrated at zero value by using blank solvents (99.9% acetone). Absorbance for chlorophylls was measured at 645 & 663nm and carotenoid at 480 & 510 nm using UV/VIS-spectrophotometer (UV-Pharma Spec 1600, Shimadzu, Japan).

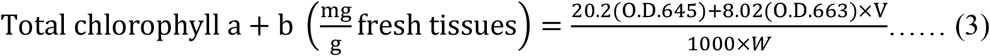

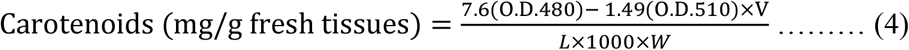

where,

V = Final volume of chlorophyll extract in 99.9% acetone.

W = Fresh weight of leaf tissue.

O.D = Optical density at a given wavelength.

L= Length of light path (1cm).

#### 2.8.2. Extraction of total phenol content

Total phytophenol was estimated in 1g of fresh leaf pieces boiled with 10 ml of 99.9% (v/v) acetone on a water bath for 10 minutes. Then, solution was allowed to centrifuge at 5000 rpm for 20 min at 25^0^C. One ml supernatant was reacted with 1ml Folins reagent and 2ml of 20% sodium carbonate to form the blue color, followed by boiling for five minutes. The final volume was adjusted up to 25ml by adding distilled water and the maximum absorbance of the blue-colored solution was read at 590 nm wavelengths.

#### 2.8.3. Estimation of rhizome pigments

Curcumin and oleoresin were extracted by dissolving 1g rhizome powder in 10 ml 99.9% v/v and kept overnight at room temperature. The filtered solution was diluted up to 10^3^ ml with acetone. The curcumin was quantified by recording absorbance reading at 425nm wavelength, while Oleoresin was detected in an air-dried solution. The oleoresin and curcumin contents were calculated using Eqs. (4) & (5) (Singh 2017; AOAC 1975):

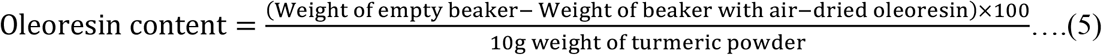

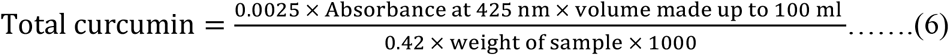

Absorbance of standard solution of curcumin 0.25g/L at 425 nm = 0.42.

### 2.9. Data Analysis

The data were statistically analyzed by applying two-way ANOVA at significant level P≤0.05 using R i386 3.4.1 and SPSS 16.0 software.

## 3. Results

### 3.1 Morphological and molecular characteristic of pathogen and biocontrol agents

#### 3.1.1 Colletotrichum gloeosporioides

Morphologically, *Colletorichum* sp. has showed abundant aerial mycelium by forming concentric pattern and ashy colony after incubation of seven days at 27±1°C. Microscopically, each conidium was observed to be in the fusiform shape. Condium has centrally placed large oil globules. The high number of sporulation in culture was recorded after 12 hours of maintenance in light and dark conditions alternatively.

#### 3.1.2 Trichoderma viride

*Trichoderma viride* was colonized up to 90mm diameter in culture plate after incubation. The colony’s color was observed fairly translucent or watery white with concentric halos. Observation with compound microscope of 100X resolution revealed microscopic structures such as frequently branched conidiophores; paired, flask-shaped phialides; and globose-shaped conidia. The opposite side of the culture media was pigmented with pale yellow color due to release of some non-volatile compounds.

#### 3.1.3 Trichoderma harzianum

The cultural plate growth of *Trichoderma harzianum* was measured 8.5cm. The colony growth was found quite slow. Initially, the aerial mycelium growth media appeared white and then acquired green, yellow shades due to abundant conidial production. Microscopically, conidiophore was observed to be branched frequently and verticillately arranged, phialides were ampuliform-convergent and conidia was sub-globous to ellipsoid in shape. The reverse side of the culture plate has exhibited intense yellow to dark orange pigmentation in media.

#### 3.1.4 Trichoderma virens

The entire plate was covered with growth and appeared as cottony, fluffy, fringed-aerial, floccose mycelium and dark green in color. Conidiophores appeared rarely branched; phialides were lageniform, convergent type; and conidia were sub cylindrical to ovoid shape.

The purification of new molecularly identified complete genomic sequences was achieved by eliminating the primer residue using Bioedit software. Thereafter, BLAST analysis has revealed 99-100% genome homology of *C. gloeosporioides, T. viride, T. harzianum* and *T. virens* with the existing database of NCBI. The confirmed nucleotide database of each isolate was registered in the Gene Bank of NCBI and the accession numbers were acquired for *C. gloeosporioides, T. viride, T. harzianum* and *T. virens*, as AMUCG1 (MK765035), AMU TVI1 (MK764992), AMU THR1 (MK765028) and AMUTVR2 (MK774725) respectively.

### 3.2. Pathogenicity test

The pathogenicity of *C. gloeosporioides* on turmeric var. sogandham was confirmed at four different doses of pathogen inoculum @ 1×10^6^, 2×10^6^, 3×10^6^ and 4×10^6^ cfu/ml, during 2015-2016. The inoculated plants of 3-4 leaf stage showed infectivity, but its severity varied according to inoculum dose, as shown in Fig.1. The spore concentrations @ 1×10^6^, 2×10^6^, and 3×10^6^ cfu/ml revealed mild severity of leaf spot disease, as characterized by numerous necrotic spots. However, the inoculum dose @ 4×10^6^ cfu/ml caused fair mortality in the form of severe drying and wilting in plants. The disease severity and leaf area of inoculated plant were calculated using Eq. (1) and (2) respectively. The morphological and microscopic characteristics of the re-isolated pathogen and the pathogen isolated from farmer’s field were observed to be identical, which satisfies the Koch’s postulate of pathogenicity.

**Fig 1.**
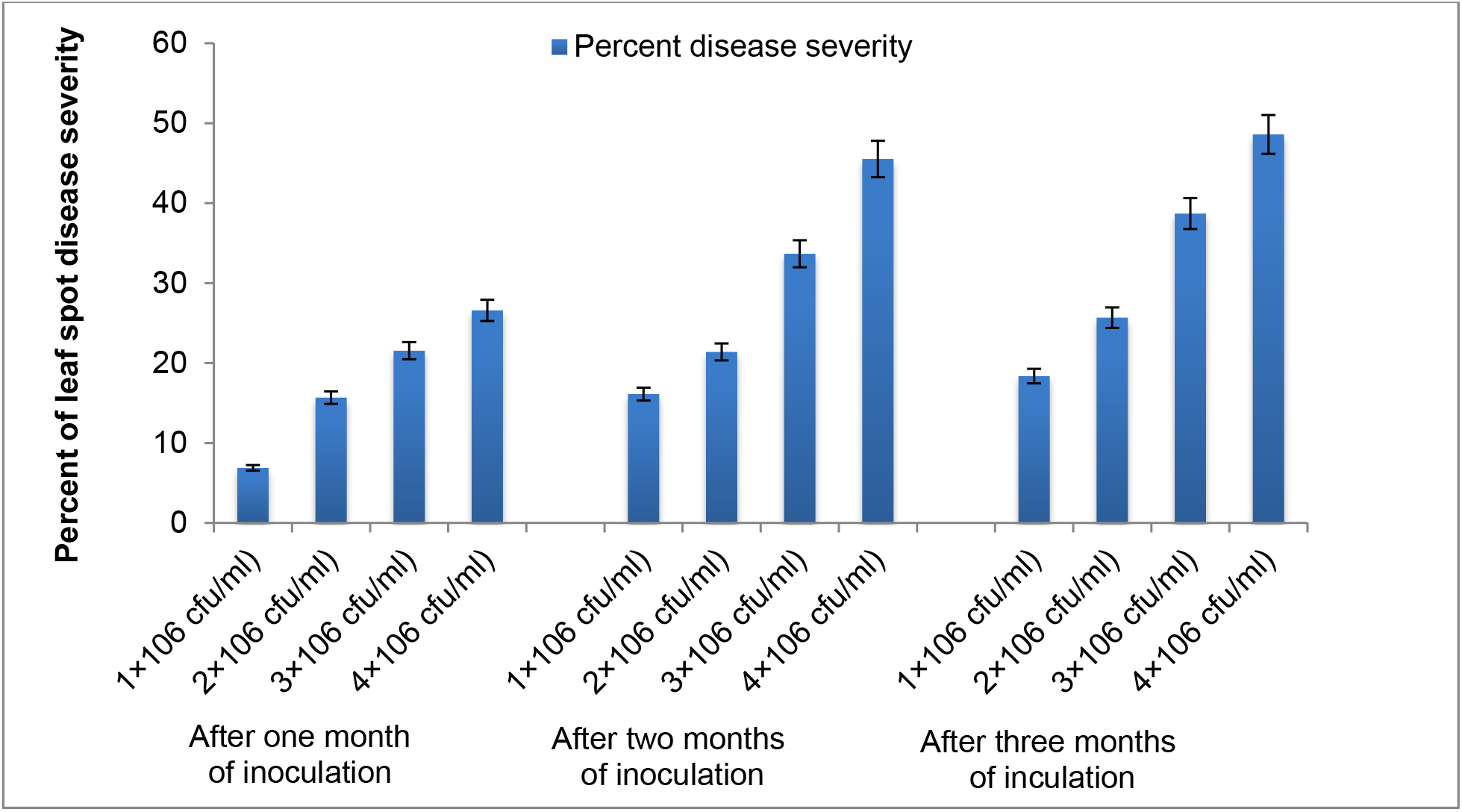
Virulence effect of *C. gloeosporioides* at different inoculum doses (2015-2016).

### 3.3. Physiological and biochemical attributes

In pot trials, the phytochemical and physiological activities of non-infected and infected plants were enhanced remarkably as presented in table 3 and table 4. The propiconazole @ RD, *A. indica* @ 50ml of 40% v/v and *T. viride* @ 50ml of 4×10^6^cfu/ml foliar sprays offered great enhancement in accessory phytochemicals concentrations: total chlorophyll (a+b)-3.78, 2.68 and 2.64 mg; carotenoid-0.9781, 0.9564 and 0.8283 mg; total phytophenol-74.74, 70.59 and 68.24µg; and oleoresin 8.585, 7.868and 7.659 %, in inoculated plants. However, improvement in curcumin contents was non-significant in each treatment (Table3). Subsequently, the physiological elements were recorded as A_N_ 0.1882, 0.1789and 0.1772 g m^-2^ day^-1^; T_N_ 3.91, 3.29 and 3.25 mmolm^-2^ s^-1^; g_s_ 3.15, 2.64 and 1.62 mmolm^-2^ s^-1^; and P_N_ 1.46, 0.9852and 0.9847 µmol m^-2^s^-1^, with high values in inoculated plants with respect to the above three treatments over control (Table 4). After one-year pot trial, the efficacy of these treatments were further noticed in the enhancement of physiological and phytochemical characteristics of naturally infected plants under open field conditions for two successive years (Table 6 and Table 7). Amongst the fungicides, biocontrol agents and phytoextracts, propiconazole @ RD, *T. viride* and *A. indica* were found most efficient to improve the essential physiological and phytochemical traits during the field trails.

The treatments have noticeably improved the plant health by increasing the quantity of chlorophyll carotenoid and phytophenol content, which is linked to trigger the defense reaction of leaf spot bio-stress in infected plants for their survival. Hence, the increased quantity of photochemicals has resulted better photosynthetic performance. Propiconazole, *A. indica* and *T. viride* sprays were found most efficient in inducing physio-biochemical tolerance against necrotic spots under both pot and field conditions. The lower disease incidence has safeguarded the proper mechanism of opening and closure of stomata, which showed good link with enhanced stomatal conductance, photon capturing in mesophyll cells, diffusion of atmospheric Co_2_ into intercellular matrixes, water balance in plants via transpiration and photosynthesis over control.

Among all the treatments, propiconazole (HRD), *A. indica* and *T. viride* were caused maximum suppression to leaf spot severity and increase the healthy leaf area of inoculated turmeric plants during 2016-2019 succeeding years of cropping seasons. In pots experiment, after sprays of propiconazole (HRD)> *T. viride* > *A. indica*, the disease incidence was recorded very low at every spray (14.76, 18.19 & 23.21%) > (18.26, 23.45 & 28.75%) > (19.16, 27.46 & 31.49%), whereas the average of leaf area index was noticed to be enhanced (2.58, 3.70 & 4.28)> (1.72, 2.93 & 3.58)> (1.59, 2.64, 3.21) in presence of pathogenic infection (Fig.2). Moreover, under field conditions foliar application of propiconazole HRD > *T. viride*> *A. indica* were also found highly effective to reduce the leaf spot severity (31.35%, 43.27% and 44.32%), and to increase leaf area index (4.63, 4.15 and 3.51) in first year trial. However, in the second year, the leaf spot severity was decreased (29.54%, 41.56% and 44.21%) but the leaf area index was improved further (4.67, 4.21 and 3.58) over control (Fig. 3). Hence, propiconazole at RD, *T. viride* and *A. indica* treatments have shown low disease intensity and maximum enhancement of physio-biochemical tolerance of infected plants compared to other treatments under both polyhouse (Fig 2) and field (Fig. 3) conditions. Thus, these treatments can be used in integrated disease management (IDM) practices for the protection of crop and improvement of crop physio-biochemistry.

**Fig 2.**
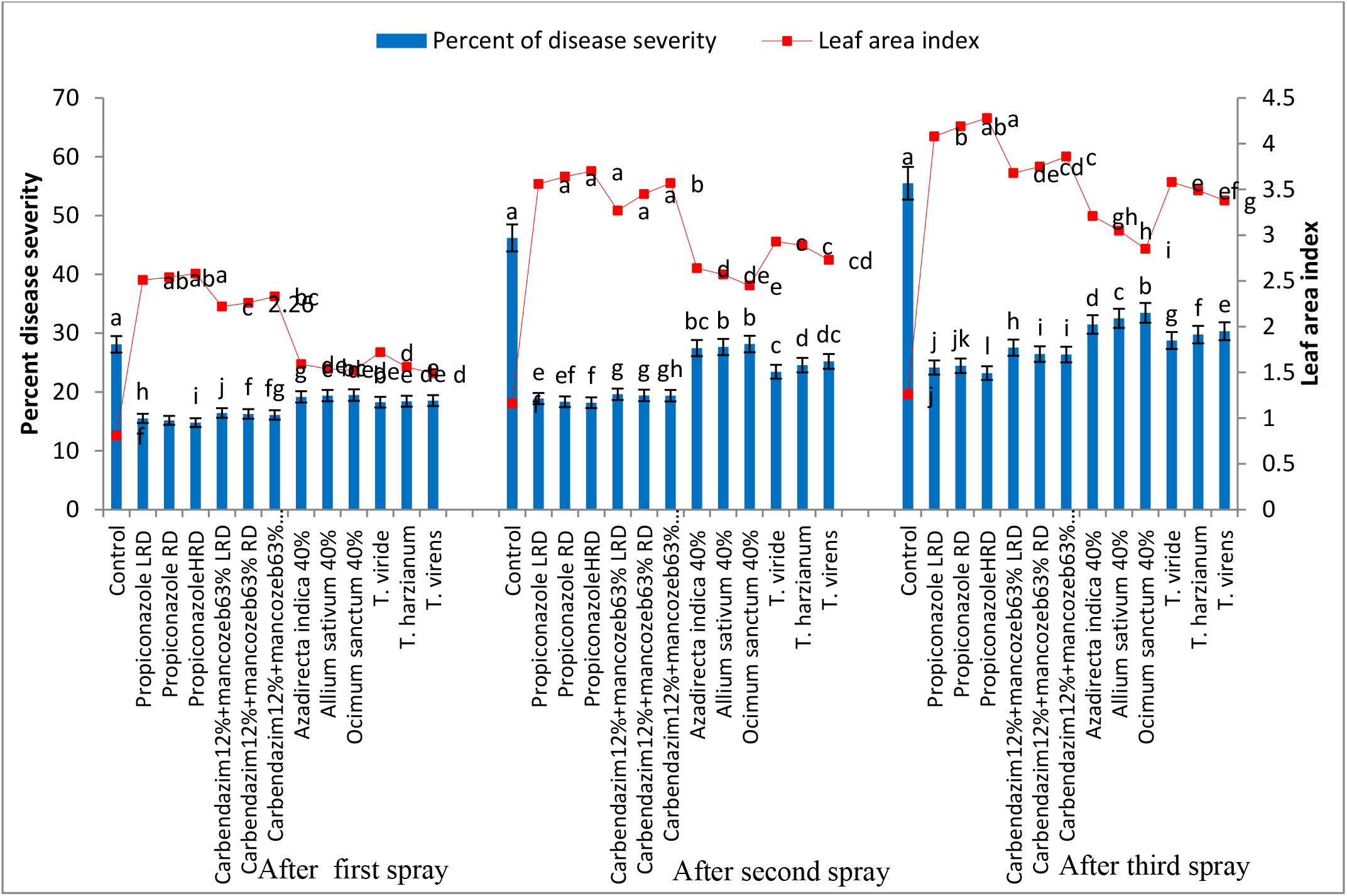
Effect of fungicides, phytoextracts and biocontrol agents foliar application on severity of leaf-spot and leaf area index of turmeric under polyhouse conditions (2016-2017).

**Fig 3.**
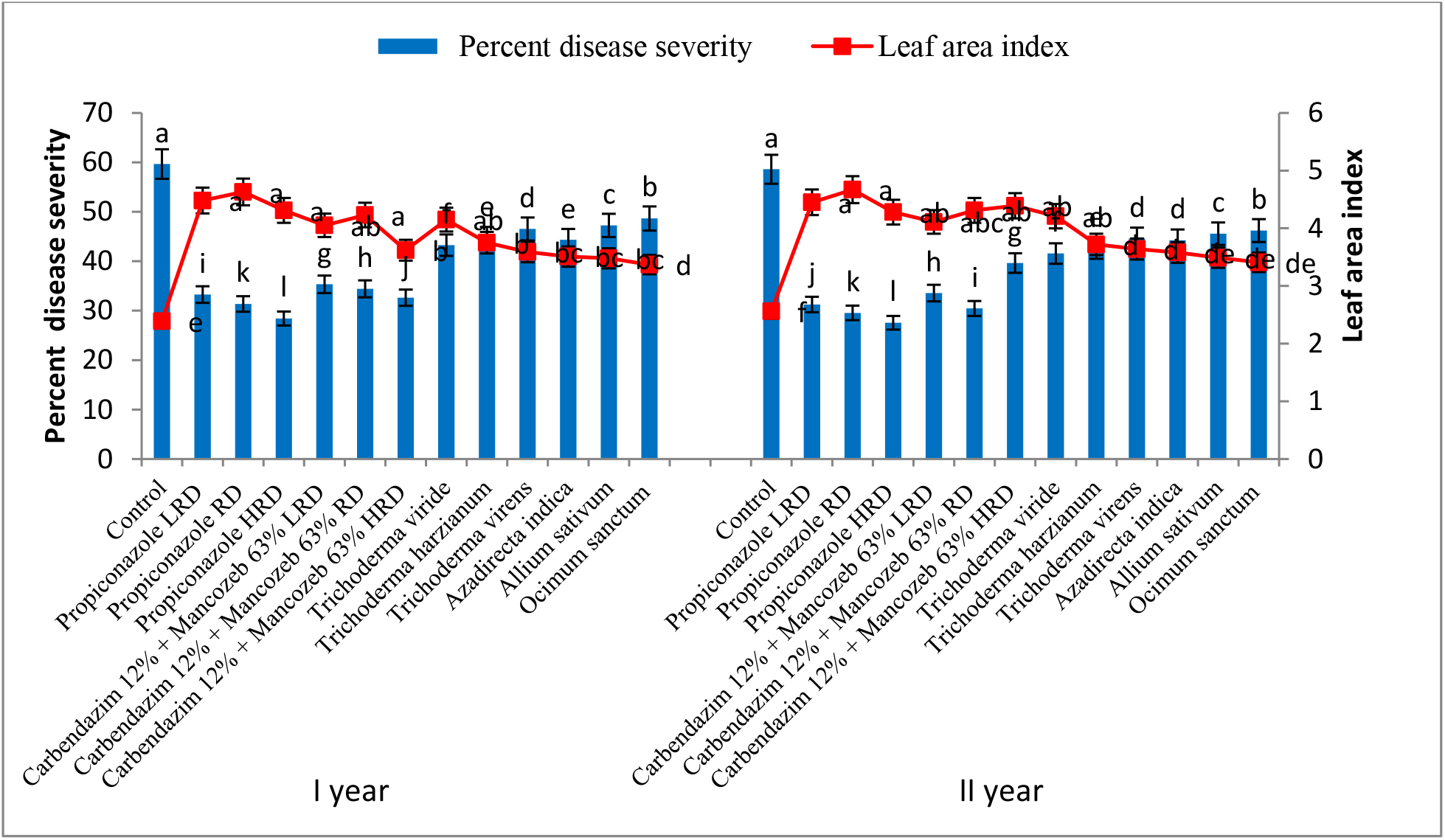
Effect of fungicides, phytoextracts and biocontrol agents foliar treatment on severity of leaf-spot and leaf area index of turmeric, under field conditions (2017-2019).

## 4. Discussion

The present study has evaluated the effects of fungicides, biocontrol agents and phytoextracts agents on the physiological and biochemical mechanisms in plants to increase their tolerance level under stress conditions of the polyhouse and field. The pathogen inoculation with micro-environment interaction could also complement the field selection under varied conditions of environment (include both pathogenic stress and environment constrains). Therefore, these treatments caused physio-biochemical variations in the treated and non-treated plants to regenerate tolerant line against pathogenic stress. Accordingly, the influence of physio-biochemical tolerance in plants can be classified into primary and secondary categories: the primary offers defense mechanisms in plants against biological and environmental stress, while the secondary improves growth characteristics to meet the demand of successful cultivation. Though many techniques have been developed to induce tolerance in plants, induction of phyto-physio-biochemical tolerance by foliar application of fungicide, biocontrol agents and phytoextracts is a new approach. The plant plasticity was modulated by the main physiological and biochemical processes to increase the plant tolerance against pathogen (Oh et al., 2009a and Oh et al., 2009b). Some plants develop thick cuticular layer in the above-ground part (stem and leaves), which leads to reduce the transpiration rate and maintain water turgor (El Ghazali 2020).

The disease density of the entire surface of turmeric leafs was recorded high under the influence of manual or natural high inoculum pressure of *Colletotrichum* sp. pathogen during polyhouse and field trials. In India, the leaf spot disease of turmeric has been found a major limiting factor in rhizome production (Kangjam et al. 2017). Therefore, the current study shows the isolation of *C. gloeosporioides* from the infected leafs of turmeric and *Trichoderma* species from turmeric’s rhizosphere. Thereafter, the pathogenicity of *C. gloeosporioides* on turmeric plants were performed at different inoculum loads under polyhouse condition. The present findings support those of Tapia-Tussell et al. (2008); Cai et al. (2009); Sekhar et al. (2017); Pasuvaraji et al. (2013); Boruah et al. (2015). While pathogenicity elicits quick and continuous changes in genes regulation in response to physio-biochemical functions, the physio-biochemical traits play a vital role in shaping the plant responses to environment. The plasticity of the plants is associated with the accumulation of bioactive molecules that increase the tolerance to stresses by modulating the main physiological and biochemical processes. The efficacy of foliar application of fungicides, biocontrol agents and phytoextracts was expanded to improve the improve the plant physio-biochemical traits of infected and non-infected turmeric plants under polyhouse conditions. Subsequently, the treatment efficiency was checked on such plants that received natural infestation of pathogen by air or soil borne inocula under field during 2017-2019. The experiment has demonstrated that the foliar application of these treatments could effectively cure the disease and strengthen the physio-biochemical mechanisms. Several researches have reported that the *Colletorichum* sp. infections on phylloplane of host plants showed negative impact on various physiological responses of plants, such as gaseous exchange, transpiration rate and photosynthesis (Kozlowski et al. 2009; Lobato et al. 2010; Guerra et al. 2014). Castrol et al. (2016) and Alves et al. (2011) reported that *Colletotrichum* sp. infected leaf was unable to carry water, solutes and other photosynthates due to the death of tissues or release of toxin. Thus, the stress of pathogenic infection has caused reduction in osmotic pressure and transpiration rate, by limiting the fixation of CO_2_ in mesophyll cells. However, the positive impact of fungicides, biocontrol agents and phytoextracts on the reduction of *C. gloeosporioides* infection and improvement of physio-biochemical characteristics on turmeric or other host plants is unique finding of the present study. Good health of plant was attributed to improved physiological characteristics, contributed by enhanced phytochemical synthesis (Dallagnol et al. 2015). The present study has demonstrated the positive impact of each treatment in the improvement of plant’s physiological response, biochemical constituents and reduction of disease rate under both polyhouse and field conditions of turmeric plants. The treatments were noted to increase the existing concentration of chlorophyll and carotenoids pigments, which have capacity to capture light in antenna complex through Photosystem II. The increase of photo-pigments led to increase the photosynthetic rate because they serve as the prime source to activate the photosynthetic gene. Nie et al. (2013) reported that higher circulation of organic carbon into roots system could raise the biomass and diameter of root. Van der Kooi et al. (2016) observed enhancement of photosynthetic machinery by the elevation of intercellular CO_2_ concentration in mesophyll cells. However, the impact of each treatment on phytophenol accumulation was significant to activate the plant defense mechanism in the presence of inoculum pressure. Thus, the plant’s phenol-content had also played great role in the productivity upgradation. Moreover, these treatments not only immune the plants for sustaining survival under stressed environmental conditions but also secure all vital machinery of plants circulating on normal path. Hence, the present treatments were proved to be promising to control the pathogenicity of *C. gloeosporioides* and to enhance the physio-biochemical traits by increasing immunity of turmeric plants against the pathogen. Jagtap et al. (2013) reported that three foliar sprays of propiconazole, *T. viride* and extracts of *Pentalonia logifolia* on turmeric plants had reduced the severity of leaf spots caused by *C. capsici* grown under pots. Yadav *et al*. (2017) also achieved the best results with foliar sprays of propiconazole and neem leaves extracts in minimizing the disease severity caused by same pathogen *C. capsici*. Currently, the fungicidal application on plants under both biotic and abiotic disease pressure was found non-acceptable due to persistence of toxic residual effects on environment or built-up resistance in pathogen over excessive application. Conversely, phytoextracts and biocontrol agents would be safer and acceptable in agricultural system for curing soil and foliar disease, besides contributing to increasing the crop productivity without any hazardous effects on plants or ecological biodiversity. The authors’ previous study (Musheer et al. 2019) determined the best result of propiconazole, *T. viride* and neem cake foliar sprays in declining the turmeric leaf spots disease incited by *C. gloeosporioides* and in enhancing the plant height, rhizome girth, fresh rhizome weight, dry rhizome weight, photopigments of leaves and curcumin content of rhizome.

## 5. Conclusion

The main aim of this study was to understand the effects of fungicides, biocontrol agents and botanical extracts on plant physiological and biochemical activities, which were noticed to be mainly associated with normal plant growth and development under the influence of pathogenic infestation. *Curcuma longa* leaf necrosis greatly affects the rhizome productivity due to death of leaf tissues; thereby plants lose normal physiological and phytochemical functioning at high rate of disease density. Therefore, suitable fungicides, botanical extracts and biocontrol agents were used on phyllosphere region of plants to minimize the severity of disease and improve the plant’s physiological machinery by enhancing the photo-pigmentation like chlorophyll and carotenoid as well as defense molecules. All treatments were found to strengthen the plant growth over control. From the trails, we concluded that propiconazole, *A. indica* and *T. viride* have high potential in managing necrotic brown blotches on leaves under both polyhouse and field conditions. Among all treatments, these three revealed the best results to improve the plant’s physio-biochemical defense mechanisms and stability of survival. Use of chemical controls to pathogen etiology is often difficult and costly, and it leads to bio-resource disintegration. Moreover, excessive dependency on synthetic chemicals for the management of pathogen can cause environmental pollution, and being non-biodegradable causes toxic residual effects in soil, water table, humans and animals. Hence, biocontrol agents and extracts of plants were added in integrated disease management modeling. However, botanicals and biocontrol extracts have showed significant results in the improvement phyto-physio-biochemical traits against *C. gloeosporioides* over control. Hence, early detection of pathogen infection and proper utilization of botanical extracts, biocontrol agents and fungicides at lower than recommended dose would be effective in inducing plant tolerance by improving physio-biochemical traits; this suppresses the disease severity before crop crosses the level of economic threshold where all disease control measures fail. This is an attractive alternative approach for the development of biotic stress tolerance in herbaceous plants. However, the mechanisms of each physio-biochemical elements require extensive study on how they respond to stresses.

## Declarations

### Ethical approval

This article does not contain any studies with human participants or animals performed by the authors.

### Consent to participate

Not applicable

### Consent for publications

Not applicable

### Availability of data and material

The genome database of isolated *Trichoderma* spp. and *Colletotrichum gloeosporioides* were deposited in National Centre of Biotechnology Information (NCBI) and acquired specific accession number of each isolate by which the nucleotide sequence records of each fungus could be generated: AMUCG1 (MK765035), AMU TVI1 (MK764992), AMU THR1 (MK765028) and AMUTVR2 (MK774725).

### Code availability

Not applicable

### Authors’ contributions

NM has special involvement in isolation, and morphological and molecular characterization of pathogen associated with turmeric leaf spot. Pot and field trials were conducted to examine the efficiency of fungicides, botanical extracts and biocontrol agents to enhance innate phyto- physiobiochemical tolerance of plants and immune the turmeric plants against *C. gloeosporioides* infection. NM, SA and AJ have approved the accuracy or integrity related to any part of the manuscript before submission.

